# Synthetic Photosynthetic Consortia Define Interactions Leading to Robustness and Photoproduction

**DOI:** 10.1101/068130

**Authors:** Stephanie G. Hays, Leo L.W. Yan, Pamela A. Silver, Daniel C. Ducat

**Affiliations:** Department of Systems Biology, Harvard Medical School, Boston, MA, USA; Wyss Institute for Biologically Inspired Engineering, Harvard University, Boston, MA, USA; MSU-DOE Plant Research Laboratory, Michigan State University, East Lansing, MI, USA; Department of Biology, Washington University in St. Louis, St. Louis, MO; Department of Biochemistry & Molecular Biology, Michigan State University, East Lansing, MI, USA.

**Keywords:** synthetic biology, photoproduction, synthetic consortia, microbial communities

## Abstract

**Background:** Microbial consortia composed of autotrophic and heterotrophic species abound in nature, yet examples of synthetic communities with mixed metabolism are limited in the laboratory. We previously engineered a model cyanobacterium, *Synechococcus elongatus* PCC 7942, to secrete the bulk of the carbon it fixes as sucrose, a carbohydrate that can be utilized by many other microbes. Here, we tested the capability of sucrose-secreting cyanobacteria to act as a flexible platform for the construction of synthetic, light-driven consortia by pairing them with three disparate heterotrophs: *Bacillus subtilis, Escherichia coli*, or *Saccharomyces cerevisiae*. The comparison of these different co-culture dyads reveals general design principles for the construction of robust autotroph/heterotroph consortia.

**Main findings:** We observed heterotrophic growth dependent upon cyanobacterial photosynthate in each co-culture pair. Furthermore, these synthetic consortia could be stabilized over the long-term (weeks to months) and both species could persist when challenged with specific perturbations. Stability and productivity of autotroph/heterotroph co-cultures was dependent on heterotroph sucrose utilization, as well as other species-independent interactions that we observed across all dyads. One interaction we observed to destabilize consortia was that non-sucrose byproducts of photosynthesis negatively impacted heterotroph growth. Conversely, inoculation of each heterotrophic species enhanced cyanobacterial growth in comparison to axenic cultures Finally, these consortia can be flexibly programmed for photoproduction of target compounds and proteins; by changing the heterotroph in co–culture to specialized strains of *B. subtilis* or *E. coli* we demonstrate production of alpha-amylase and polyhydroxybutyrate, respectively.

**Conclusions:** Enabled by the unprecedented flexibility of this consortia design, we uncover species-independent design principles that influence cyanobacteria/heterotroph consortia robustness. The modular nature of these communities and their unusual robustness exhibits promise as a platform for highly-versatile photoproduction strategies that capitalize on multi-species interactions and could be utilized as a tool for the study of nascent symbioses. Further consortia improvements via engineered interventions beyond those we show here (i.e. increased efficiency growing on sucrose) could improve these communities as production platforms.

## BACKGROUND

Cyanobacteria are under increased investigation as alternative crop species for the production of fuels and other commodity chemicals. While much research focuses on engineering cyanobacterial metabolism towards the synthesis of end products (e.g. biofuels), cyanobacteria are also under consideration for the production of carbohydrate feedstocks to support fermentative bioindustrial processes [1–3]. In this approach, cyanobacterial biomass is processed to provide [4–6] or manipulated to secrete simple fermentable sugars [2, 3, 7–10]. Multiple groups have recently reported that different cyanobacterial species are capable of secreting soluble sugars in considerable quantities, and continuous production can be sustained over long time periods. The specific productivities of these strains are high in comparison to plant-based feedstocks [10, 11], and, additionally, cyanobacterial cultivation can use landmass and water that is not suitable for standard agriculture. The promise of this approach has led to investment by biotechnology firms to construct compatible reactors and conduct scaled pilot trials [12]. Yet, as with most cyanobacterial and algal processes, there are barriers to scaled cultivation that may make such strategies economically non-competitive [13–15]. These include the costs to recover dissolved sugars or transport culture media to fermenters as well as potential loss to contaminants. These costs might be avoided by simultaneous conversion of the cyanobacterial feedstock into higher-value compounds by co-existing microbes in a “one-pot” reaction [16].

We test this premise by characterizing a series of synthetic co-cultures in which engineered cyanobacteria fix and secrete carbon to support growth of a broad range of evolutionarily-unrelated model heterotrophs. To provide organic carbon to engineered consortia, we use a *Synechococcus elongatus* PCC 7942 strain previously engineered to export up to 85% of the carbon it fixes in the form of sucrose [10], a simple sugar also produced by plant-based feedstocks (e.g. sugarcane) that is readily consumed by many microbes. *S. elongatus* naturally accumulates sucrose as a compatible solute [1, 17], and can export this carbohydrate through heterologous expression of the proton/sucrose symporter, *cscB* (hereafter *cscB^+^*)[10, 18]. Under mild osmotic shock and slightly alkaline conditions, *cscB^+^ S. elongatus* continuously generate sucrose at up to 36mg sucrose L^−1^ hr^−1^, a rate predicted to substantially exceed that of sugarcane, if successfully cultivated to scale [10]. This strain therefore demonstrates great promise as the basis for flexible photoproduction of distinct target compounds, especially if the excreted sugars are efficiently harnessed to promote growth of a variety of useful microbes while minimizing additional processing steps.

Our approach to construct cyanobacteria/heterotroph consortia utilizes an inherently modular design: i) the “autotrophic module” (*cscB^+^ S. elongatus*) fixes carbon and secretes sucrose; while ii) a variety of species (i.e. the “heterotrophic module”) are examined for growth and productivity in pairwise co-cultures (Fig. 1A). Specifically, we examine a number of workhorse model microbes (*Escherichia coli*, *Bacillus subtilis*, or *Saccharomyces cerevisiae*) for their capacity to persist and grow using only cyanobacterially-derived organic products. We chose these species because they are well-researched, phylogenetically well-distributed (i.e. gram negative, gram positive, or eukaryotic), possess excellent genetic toolkits, a variety of engineered and mutant lines are readily available, and are frequently researched for bioindustrial applications. Finally, these species are not isolated from environments where cyanobacteria dominate, therefore it is expected that no pre-existing evolutionary relationship to cyanobacteria exists.

**Fig. 1.**
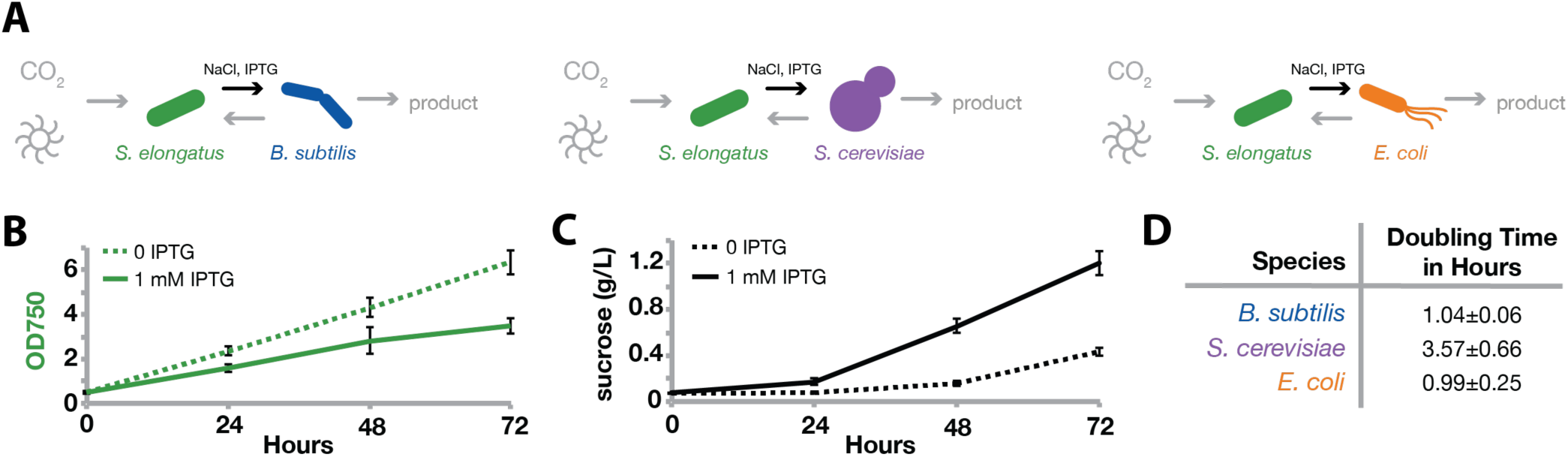
Axenic characterizations of candidate strains. (A) This schematic shows the engineered microbial community design. *CscB*^*+*^ *S. elongatus* (green) capture light and CO_2_ via photosynthesis. Fixed carbon is secreted as sucrose (black arrows) when induced with IPTG in the presence of osmotic pressure (NaCl). This secreted carbon then supports the growth of *B. subtilis* (blue), *S. cerevisiae* (purple), or *E. coli* (orange) with the final goal of production of target compounds from those heterotrophs. Axenic *cscB*^*+*^ *S. elongatus* was grown in ^CoB^ BG-11 with (solid line) and without IPTG (dashed line) to induce sucrose secretion. Cell density (B) and sucrose levels in culture supernatants (C) were measured. Error bars are standard deviation of 8 biological replicates. For characterization of cyanobacteria in ^CoY^BG-11 see Additional File 1: Fig. S1. (D) Heterotroph growth in isolation was characterized via growth rate in co-culture buffer supplemented with 2% sucrose. Error is standard deviation of ≥ 3 replicates.

Taken together, this modular framework permits us to survey the capacity of the *cscB*^*+*^ *S. elongatus* strain for construction of diverse artificial consortia with distinct composition and functions, as well as permits between-species comparison of consortia features. For example, there may be constraints to coexistence of cyanobacteria and heterotrophs that are shared across different microbial species and would be expected impair co-culture stability. By identifying such bottlenecks, it is then possible to optimize the system for improved performance and bioproduction through strain engineering or culturing techniques.

In this work, we demonstrate that *cscB*^*+*^ *S. elongatus* can promote growth of all tested heterotrophic species in pairwise co-cultures without the addition of external carbohydrates. We identify non-programmed interactions between cyanobacteria and heterotrophic microbes, including phenomena that are shared across all species. In some cases, these emergent interactions limit heterotrophic growth and/or stability. By mitigating bottlenecks to heterotrophic growth, we demonstrate that each of our co-cultures can persist over the long-term (days to months): an important feature that is often missing in synthetic cross-feeding consortia [19–22]. We also observe that the presence of each tested heterotroph species promotes the growth of *S. elongatus*, despite a lack of pre-existing evolutionary relationships. The modular design of composition allows these consortia to be flexibly functionalized for photoproduction of a target metabolite (polyhydroxybutyrate, PHB) or a target enzyme (alpha amylase) with appropriate *E. coli* and *B. subtilis* strains, respectively. Finally, we examine our results for general design principles for improvement of engineered cyanobacteria/heterotroph consortia, and discuss potential of this platform for the “bottom-up” assessment of naturally-occurring microbial communities.

## RESULTS

### Cyanobacteria in consortia with heterotrophs

We designed pairwise consortia where *cscB^+^ S. elongatus* secrete sucrose in response to osmotic pressure and isopropyl β-D-1-thiogalactopyranoside (IPTG)-induced *cscB* expression [10]. Carbon secreted by cyanobacteria promotes growth of co-cultured heterotrophs (Fig. 1A). Media with optimized compositions of nitrogen, salt, and buffer were developed: termed ^CoB^BG-11 for use in cyanobacteria/bacteria consortia and ^CoY^BG-11 for cyanobacteria/yeast co-culture (see Materials and Methods). We verified that *S. elongatus* grows and produces sucrose in both ^CoB^BG-11 and ^CoY^BG-11 (Fig. 1B, Additional File 1: Fig. S1). As previously reported [10], induction of *cscB* greatly enhances the rate of sucrose export, and this redirection of carbon resources leads to slower growth (Fig. 1B&C, Additional File 1: Fig. S1). We also verified that all heterotrophs are capable of growth when grown as monocultures in these defined media when provided with exogenous sucrose (2%) as the sole carbon source (Fig. 1D).

*CscB^+^ S. elongatus* directly support heterotroph growth in co-cultures that contain no external carbon sources (Fig. 2, Additional File 1: Fig. S2). In all consortia, *cscB^+^ S. elongatus* is inoculated with a heterotrophic microbe in the appropriate co-culture media (with or without the addition of 1mM IPTG to induce *cscB* expression; see Materials and Methods) and grown over 48 hours in constant light. We tracked the growth of *cscB^+^ S. elongatus* in co-culture through the use of flow cytometry. Viable heterotrophs were tracked by analyzing the number of colony forming units (CFUs) when plated on the appropriate solid media. More than one strain of *E. coli* and *S. cerevisiae* were analyzed in co-culture to determine the effects of a particular genetic background on growth kinetics.

**Fig. 2.**
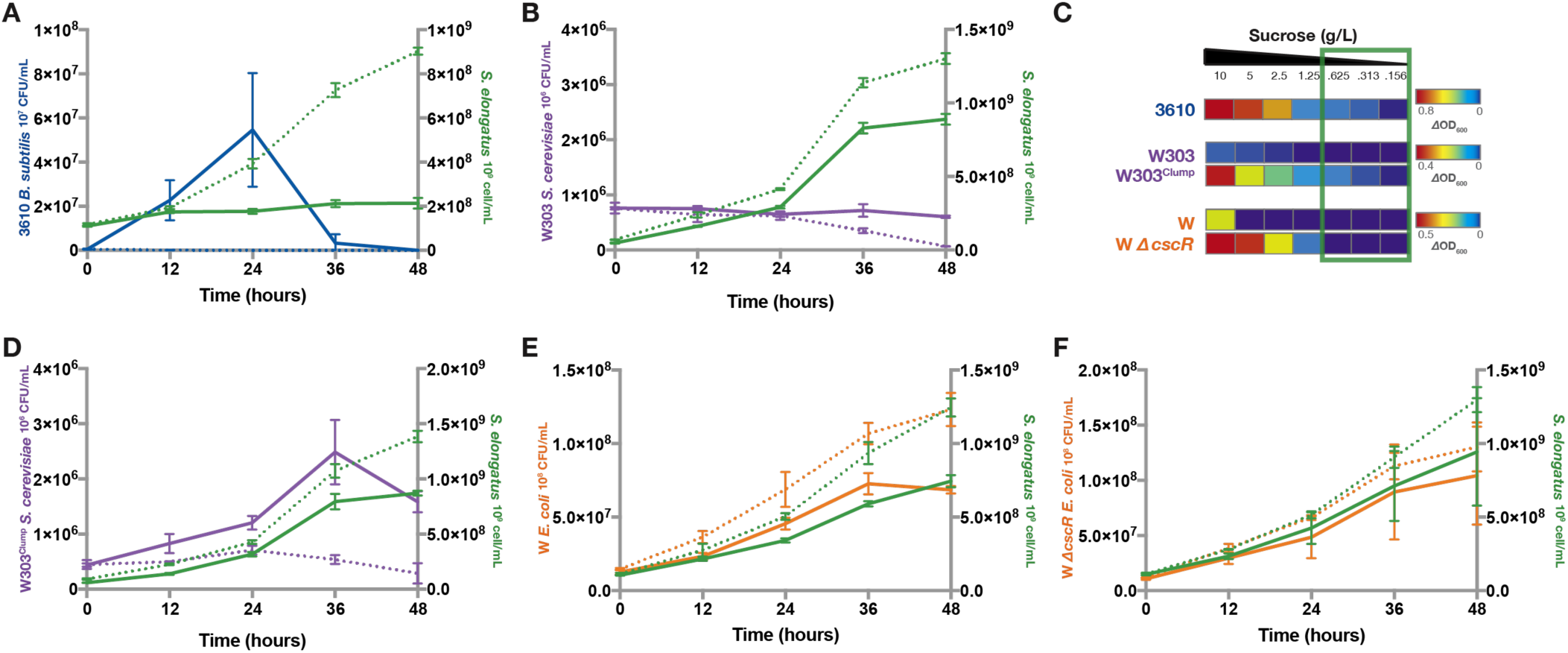
S. elongatus supports microbial communities in batch culture. Batch cultures of *cscB*^*+*^ *S. elongatus* (green) in co-culture with *B. subtilis* (blue), *S. cerevisiae* (purple), or *E. coli* (orange) were grown in constant light. *cscB*^*+*^ *S. elongatus* cells/mL were determined by flow cytometry every 12 hours for co-cultures containing *B. subtilis* (A; green), *S. cerevisiae* (B, D; green), and *E. coli* (E, F; green). Co-cultures with uninduced (dashed lines) or induced *cscB* expression (solid lines) were tested. Heterotroph viability was monitored by colony forming unit (CFU) for all *B. subtilis* (A; blue), *S. cerevisiae* (E strain W303, F strain W303^Clump^; purple), and *E. coli* (E strain W, F strain W Δ*cscR*; orange) co-cultures. Data for A, B, D, E, and F, are representative, same-day experiments where error bars are the standard error in 3 biological replicates. Additional replicates in Additional File 1: Fig. S2. (C) Axenic heterotroph growth was tested in defined media with varying concentrations; the range of sucrose that *cscB*^*+*^ *S. elongatus* can secrete in 48 hours is denoted by a green box. Average OD_600_ is shown as a metric of growth for ≥ 6 biological replicates. OD_600_ was correlated to viable colony forming units (CFU) in Additional File 1: Fig. S3. No contaminants/heterotrophic colonies grew from axenic cyanobacteria controls.

*B. subtilis* growth in co-culture is dependent on IPTG-induced sucrose secretion from *cscB^+^ S. elongatus* (Fig. 2A, Additional File 1: Fig. S2A). Without induction of *cscB* to enable sucrose secretion, *B. subtilis* fails to grow, indicating that sucrose availability is limiting at basal levels of sucrose export. However, when IPTG is added to increase sucrose export, *B. subtilis* growth is nonmonotonic: after an initial increase viability decreases during the second 24 hours of co-culture (Fig. 2A).

*S. cerevisiae* growth in co-culture is dependent on genetic engineering to improve sucrose utilization. Wild type (WT) *S. cerevisiae* W303 did not grow in co-culture with or without IPTG induction (Fig. 2B). We examined the capacity of WT *S. cerevisiae* W303 to grow axenically at low sucrose concentrations and saw poor/no growth below 2.5g/L sucrose (Fig. 2C), a higher concentration sucrose than is produced by *cscB^+^ S. elongatus* at 48 hours (Fig. 1C). We then turned to an engineered strain, hereafter referred to as W303^Clump^, derived from previous directed evolution experiments of *S. cerevisiae* W303 in low sucrose media [23]. W303^Clump^ (originally called Recreated02 in Koschwanez *et al.* 2013) contains mutations in genes CSE2, IRA1, MTH1, and UBR1 that enhance fitness in dilute sucrose, and also contains a nonsense mutation in ACE2 that compromises the full septation of budding daughter cells from the mother, resulting in small clonal cell aggregates (~6.6 cells per clump on average). These aggregates grow in low sucrose due to increased local cell concentration and increased hexose availability after extracellular cleavage of sucrose by an invertase [24]. Unlike the parental strain, axenic cultures of W303^Clump^ exhibited some growth at all tested sucrose concentrations ≥0.156g/L (Fig. 2C), as well as when co-cultured with IPTG-induced *cscB^+^ S. elongatus* (Fig. 2D). Similar to *B. subtilis*, in co-culture W303^Clump^ *S. cerevisiae* demonstrate declining viability after an initial period of growth (Fig 2D).

Finally, *E. coli* W grows in co-culture independently of induced sucrose secretion from *cscB^+^ S. elongatus* (Fig. 2B, Additional File 1: Fig. S2B&C). In axenic culture, WT *E. coli* W exhibits growth only when supplemented with >5g/L sucrose (Fig. 2C), well above the levels *cscB^+^ S. elongatus* secrete during 48 hours of growth (Fig. 1C). We therefore tested the growth of an *E. coli* W strain engineered for growth on sucrose. In this strain the sucrose catabolism repressor, *cscR,* was deleted (the strain is hereafter referred to as *ΔcscR E. coli*), resulting in more rapid growth at lower sucrose concentrations [25–27]. Indeed, monocultures of *ΔcscR E. coli* exhibit the capacity to grow on lower concentrations of sucrose (≥1.25g/L; Fig. 2C), yet they still demonstrate no growth at sucrose concentrations in the range that *cscB^+^ S. elongatus* can secrete in 48 hours (Fig. 1C, green box Fig. 2C). In co-culture, *ΔcscR E. coli* exhibited the same monotonic growth pattern as the unmodified strain (Fig. 2A&C, Additional File 1: Fig. S2B&C). This suggests that in the first days of co-culture, while exported sucrose concentrations are low (≤1g/L), *E. coli* strains cannot utilize sucrose effectively and dominantly depend on other metabolites from *S. elongatus*; perhaps extracellular polymeric substances [28, 29].

### Light driven cyanobacterial metabolism inhibits heterotroph viability

Cyanobacterial light driven metabolism is the source of heterotroph growth inhibition when sucrose is not limiting (Fig. 3A). The lack of monotonic growth in *B. subtilis* and *S. cerevisiae* co-cultures indicate that interactions beyond sucrose feeding are occurring between heterotrophs and cyanobacteria (Fig. 2A, 2F). To focus on products other than fixed carbon that influence heterotrophic viability and eliminate the confounding factor that cyanobacteria only generate sucrose in the light [10], co-cultures were supplemented with exogenous sucrose and cultivated in the light or dark. After 12 hours of co-cultivation, heterotroph viability of each of the three species was determined, revealing decreased growth or death correlated with increasing concentrations of cyanobacteria solely in illuminated cultures (Fig. 3B-D, Additional File 1: Fig. S4). This effect is most apparent in strains of *B. subtilis* (Fig. 3B, Additional File 1: Fig. S4A) where the viability of heterotrophic species decreases by orders of magnitude when co-cultured in the light with high concentrations of *S. elongatus*.

**Fig. 3.**
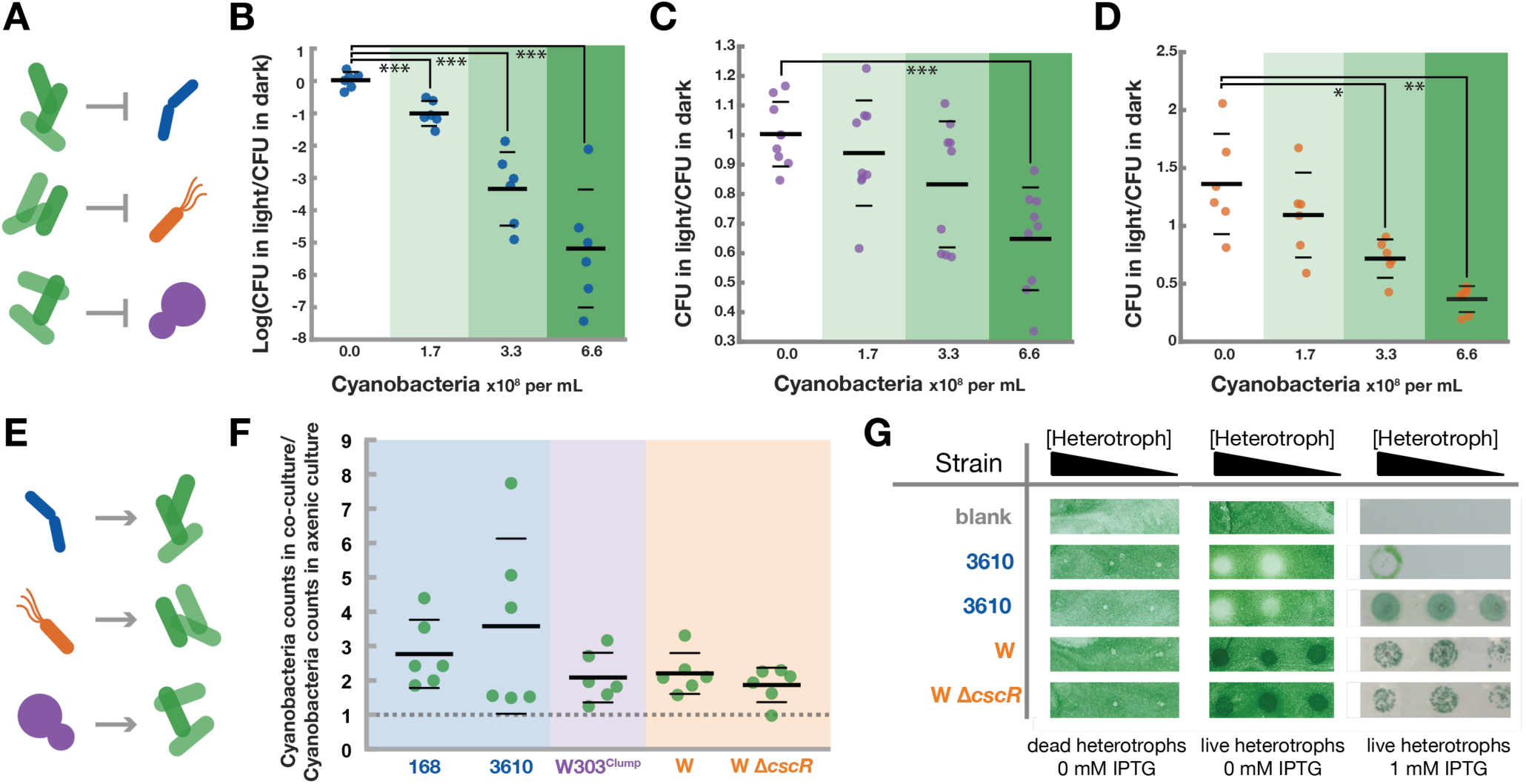
Microbial interactions. Engineered consortia demonstrate un-engineered interactions that can be classified into two categories: negative effects that cyanobacteria have on heterotrophs (A) and positive effects heterotrophs have on cyanobacteria (E). *B. subtilis* 3610 (B), W303^Clump^ *S. cerevisiae* (C), and *E. coli* W Δ*cscR* (D) were co-cultured with various concentrations of *S. elongatus* and heterotroph CFU/mL were determined after 12 hours of cultivation in either light or dark. Ratios of CFU in light compared to CFU in dark are reported (B-D). Additional strains were tested in Additional File 1: Fig. S4. P-values of two-tailed t-tests with Welch’s correction are denoted with asterisks: * 0.01 to 0.05, ** 0.001 to 0.01, *** 0.0001 to 0.001, **** < 0.0001. Positive effects of heterotrophs on cyanobacteria (E) were observed in liquid (F), evidenced by the number of cyanobacteria cells measured in co-cultures relative to axenic controls after 48 hours in constant light. These co-cultures were inoculated with two orders of magnitude fewer *cscB*^*+*^ *S. elongatus* (~1.7x10^6^cells/mL) than the co-cultures depicted in Fig. 2 (~1.7x10^8^cells/mL), and 1mM IPTG was added to all cultures to induce sucrose export. Thick horizontal lines represent the average measurement for each condition while thin horizontal lines represent one standard deviation from the mean. Positive effects of heterotrophs on cyanobacteria in previous liquid batch experiments is summarized in Additional File 1: Fig. S5. The influence of heterotrophs on cyanobacterial growth on solid media (G) was determined by plating a dilute lawn of *cscB*^*+*^ *S. elongatus* on ^CoB^BG-11 agar plates. The cyanobacterial lawn was overlaid with the specified strain in ten-fold serial dilutions of heterotroph and in constant light with or without IPTG.

### Heterotrophic species stimulate cyanobacterial growth

Conversely, co-culture with heterotrophs can stimulate growth of *S. elongatus* (Fig. 3E, Additional File 1: Fig. S5). This was observed during batch cultures when cyanobacteria counts were higher in co-cultures then in control monocultures of cyanobacteria at various time points (Additional File 1: Fig. S5). Because batch cultures of relatively dense *S. elongatus* can negatively impact heterotrophic viability (Fig. 2A & 2E, 3A-D), and also lead to significant self-shading, we inoculated low concentrations of *cscB^+^ S. elongatus* induced with IPTG in co-culture with heterotrophs. After 48 hours of co-culture, cyanobacteria numbers in co-culture were normalized to axenic cyanobacteria controls. We observe significant increases in cyanobacterial growth in the presence of heterotrophic microbes, with total cyanobacterial cell counts increasing by between 80 and 250% on average (Fig. 3F).

The growth-promoting effect of heterotrophs on cyanobacteria persists when co-cultivated on solid media. We spotted dilutions of *B. subtilis* or *E. coli* on a lawn of dilute cyanobacteria with or without IPTG (Fig. 3G). Areas of the cyanobacterial lawn overlaid with *E. coli* exhibited more rapid growth than the surrounding lawn of *S. elongatus* alone. The growth-promoting effect of *B. subtilis* on *cscB^+^ S. elongatus* was dependent upon induction of sucrose export. Without IPTG, spots of *B. subtilis* inhibited cyanobacterial growth, however in the presence of IPTG, *B. subtilis* could stimulate growth on or in the vicinity of the spot it was plated (Fig. 3G). *S. cerevisiae* was not assayed in this manner because of poor growth of *cscB^+^ S. elongatus* on ^CoY^BG-11 solid agar plates. Collectively, these experiments indicate that all three evolutionarily unrelated heterotrophs can significantly increase cyanobacterial growth under a range of growth conditions.

### Robustness in designed photosynthetic consortia

As the inhibitory effects of cyanobacteria on the viability of heterotrophs are only observed at relatively high density and in the light (Fig. 3A-D), we evaluated co-cultivation strategies that would prevent overgrowth of cyanobacteria to determine if heterotrophic viability could be maintained in the long-term. We first turned to Phenometrics environmental Photobioreactors (ePBRs), which have turbidostat capabilities in addition to control of light, temperature, and culture stirring [30].

When cultivated at a constant density, *E. coli/cscB^+^ S. elongatus* co-cultures persist over time and heterotrophic viability is maintained. Co-cultures of induced *cscB^+^ S. elongatus* and W *ΔcscR E. coli* were grown continuously in ePBRs under constant light (Fig. 4A). By monitoring *cscB^+^ S. elongatus* cell counts and *E. coli* CFUs, we determined that co-cultures maintain stable ratios for more than two weeks (Fig. 4A, Additional File 1: Fig. S6).

**Fig. 4.**
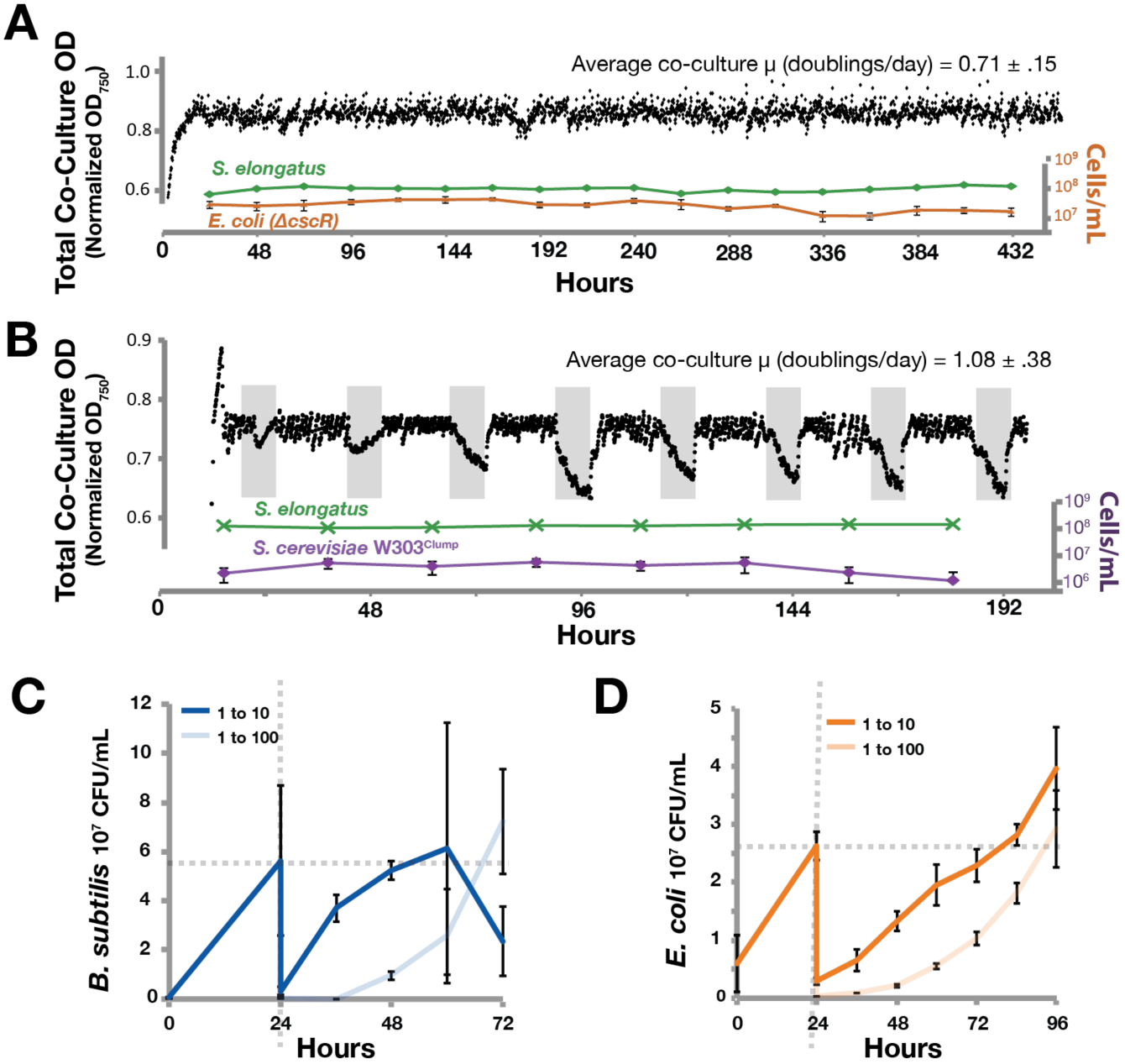
Co-cultures persist through time and perturbation. Representative continuous co-cultures of *E. coli* W Δ*cscR/cscB*^*+*^ *S. elongatus* (A) and W303^Clump^ *S. cerevisiae*/*cscB*^*+*^ *S. elongatus* (B) were cultured in photobioreactors with 1mM IPTG. *E. coli*-containing consortia were grown in constant light while *S. cerevisiae* communities were exposed to 16:8 hour light/dark photo periods (grey spaces represent darkness). Optical density of the entire culture (black points) as well as counts for the individual cell types were tracked (green *S. elongatus,* orange *E. coli* W Δ*cscR*, purple W303^Clump^ *S. cerevisiae*). Additional photobioreactor cultures for *E. coli* W Δ*cscR* and W303^Clump^ *S. cerevisiae* are presented in Additional File 1: Fig. S6 and S7, respectively. Extended W303^Clump^ *S. cerevisiae*/*cscB*^*+*^ *S. elongatus* co-cultures are presented in Additional File 1: Fig. S8. Recovery of heterotrophs in batch co-cultures of *B subtilis* 3610/*cscB*^*+*^ *S. elongatus* (C) or *E. coli* W Δ*cscR* /*cscB*^*+*^ *S. elongatus* (D) following dilution (vertical dashed grey line) was monitored by viable colony counts. Perturbations on to solid media are presented in Additional File 1: Fig. S9.

Similarly, when cultured in ePBRs, *S. cerevisiae* W303^Clump^ maintained viability in co-culture with *cscB^+^ S. elongatus* and persisted through variable light conditions. We induced *cscB^+^ S. elongatus* to secrete sucrose and inoculated *S. cerevisiae* W303^Clump^ into ePBRs programmed with an alternating diurnal illumination regime (16 hours light:8 hours dark, Fig. 4B). Sustained growth in these continuous cultures indicates that yeast persist through periods of darkness when cyanobacteria are unable to supply sucrose or other photosynthates (Fig. 4D, Additional File 1: Fig. S7). In similar experiments that were extended over longer time periods, *S. cerevisiae* W303^Clump^ maintains viability in continuous culture with sucrose-secreting *S. elongatus* for greater than two months (Additional File 1: Fig. S8).

Prokaryotic co-cultures with cyanobacteria persist through population bottlenecks and changes in environmental structure. *B. subtilis*/*cscB^+^ S. elongatus* and W *ΔcscR E. coli/cscB^+^ S. elongatus* co-cultures were subjected to large dilutions (1 to 10 or 1 to 100) to determine viability following the introduction of a population bottleneck. Cyanobacterial growth was consistently visually observed following dilution, while heterotroph growth was measured via CFU. In perturbed cultures, heterotrophs can return to pre-dilution levels within three days (Fig. 4C & 4D). We also examined the persistence of heterotrophs in co-culture following plating onto solid media after growth in liquid co-culture. Co-cultures containing *cscB*^+^ *S. elongatus* and *B. subtilis* 3610 or W *ΔcscR E. coli* were moved from liquid to solid environments and back again (Additional File 1: Fig. S9). This transfer is expected to disrupt the ratio of different species within co-culture and alters any interactions dependent on the co-culture constituents being in well-mixed environments. After incubating co-cultures in the light on agar plates that have no added carbon, green growths from the agar plate were picked into liquid ^CoB^BG-11. Picked co-cultures were recovered in constant light for 2-5 days; wells with cyanobacterial growth were determined qualitatively by presence of green coloration, and heterotroph persistence was evaluated by plating onto rich media. In the majority of cultures both cyanobacteria and the corresponding heterotroph persisted through these perturbations, although *B. subtilis* was lost from the co-culture somewhat more frequently than *E. coli* (Additional File 1: Fig. S9).

### Bioproduction from functionalized co-cultures

As multiple species can be co-cultured with *cscB^+^ S. elongatus*, it is possible to exchange heterotrophs to functionalize consortia for desired activity. In this design, the heterotrophic species of the consortia acts as a “conversion module” to metabolize the products of photosynthesis into target bioproducts in a “one-pot reaction” (Fig. 5A). We tested two heterotrophic strains capable of producing distinct products: enzymes (Fig. 5B&C) and chemical precursors (Fig. 5D).

**Fig. 5.**
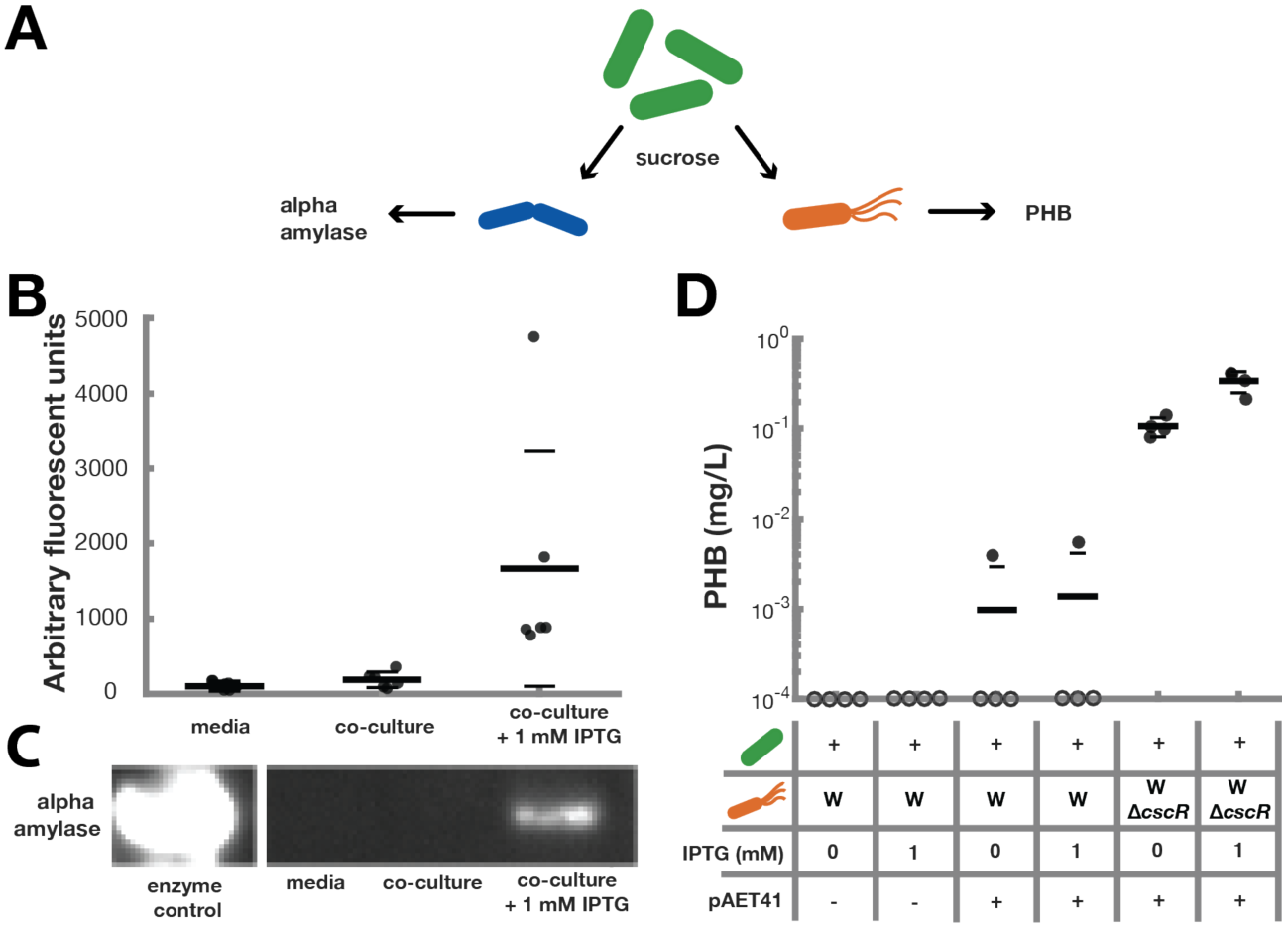
Photoproduction of enzymes and metabolites from co-culture. Flexible functionalization of co-cultures was accomplished via the addition of heterotrophs capable of producing target compounds (A). Alpha-amylase is naturally produced and secreted by *B. subtilis* strain 168. Supernatants from 24 hour cultures of *B. subtilis* 168 alone or in co-culture with *cscB^+^ S. elongatus* were tested for enzymatic activity (B). Western blots also reveal the presence of alpha-amylase in co-cultures containing IPTG (C). *E. coli* is capable of making PHB when carrying the pAET41 plasmid. Batch co-cultures of *E. coli* (with or without pAET41 to enable PHB production) and *cscB*^*+*^ *S. elongatus* were cultivated for one week with or without IPTG to induce sugar. PHB content of the total culture was analyzed (D). Filled circles represent measured values; hollow circles placed on the x-axis represent cultures in which no PHB was formed or was produced at levels below the detection limit. Thick horizontal lines represent the average measurement for each condition while thin horizontal lines represent one standard deviation from the mean.

Alpha-amylase is produced in co-cultures of *B. subtilis* strain 168 and *cscB^+^ S. elongatus*. *B. subtilis* is a chassis for enzyme production [31] and strain 168 naturally produces active alpha-amylase [32]. In consortia with *S. elongatus*, *B. subtilis* 168 produces alpha-amylase after 24 hours in constant light (Fig. 5A&B). The resulting alpha-amylase is functional as determined by enzymatic assay, and accumulates at significantly higher levels in co-cultures with *cscB^+^ S. elongatus* induced to secrete sucrose (Fig. 5A) in comparison to other co-cultures.

Engineered *E. coli/S. elongatus* communities produce PHB. We co-cultured *E. coli* strains harboring a previously described PHB production plasmid, pAET41 [33] with *cscB^+^ S. elongatus* for one week in constant light and measured PHB in the total biomass by liquid chromatagraphy (Fig. 5D). While production from the WT *E. coli* W strain is similar with and without IPTG, the *ΔcscR E. coli* W mutants that utilize sucrose more effectively produce significantly more PHB (Fig. 2C, Fig. 5D)[27]. Furthermore, upon the addition of IPTG, the *ΔcscR E. coli* W strain can produce three times as much PHB in co-culture than in un-induced consortia. Taken together, these results demonstrate that consortia can be flexibly programmed for photoproduction of different bioproducts by employing different heterotrophic organisms.

### Mathematical framework describing interactions

We summarized the interactions that were deliberately programmed in our synthetic co-cultures into a simple mathematical framework, illustrated in Fig. 6A and described in the Supplemental Material (Additional File 1: Mathematical Framework). Briefly, the population of phototrophs, *P(t),* at time *t* can be approximated with a linear growth rate of *μ*_*p*_, based on the experimental data in Fig. 1B and 1C. The sucrose concentration, *S(t)*, is dependent both upon the production from cyanobacteria, *α*, and the consumption of sucrose by the heterotrophs, *β*. The heterotroph population, *H(t)*, is dependent on the maximal growth rate of each heterotroph on sucrose (μ_max_), the sucrose concentration at which the growth rate is half maximal (*S*_*halfmax*_), and the sucrose concentration (*S(t)*)[34, 35].

**Fig. 6.**
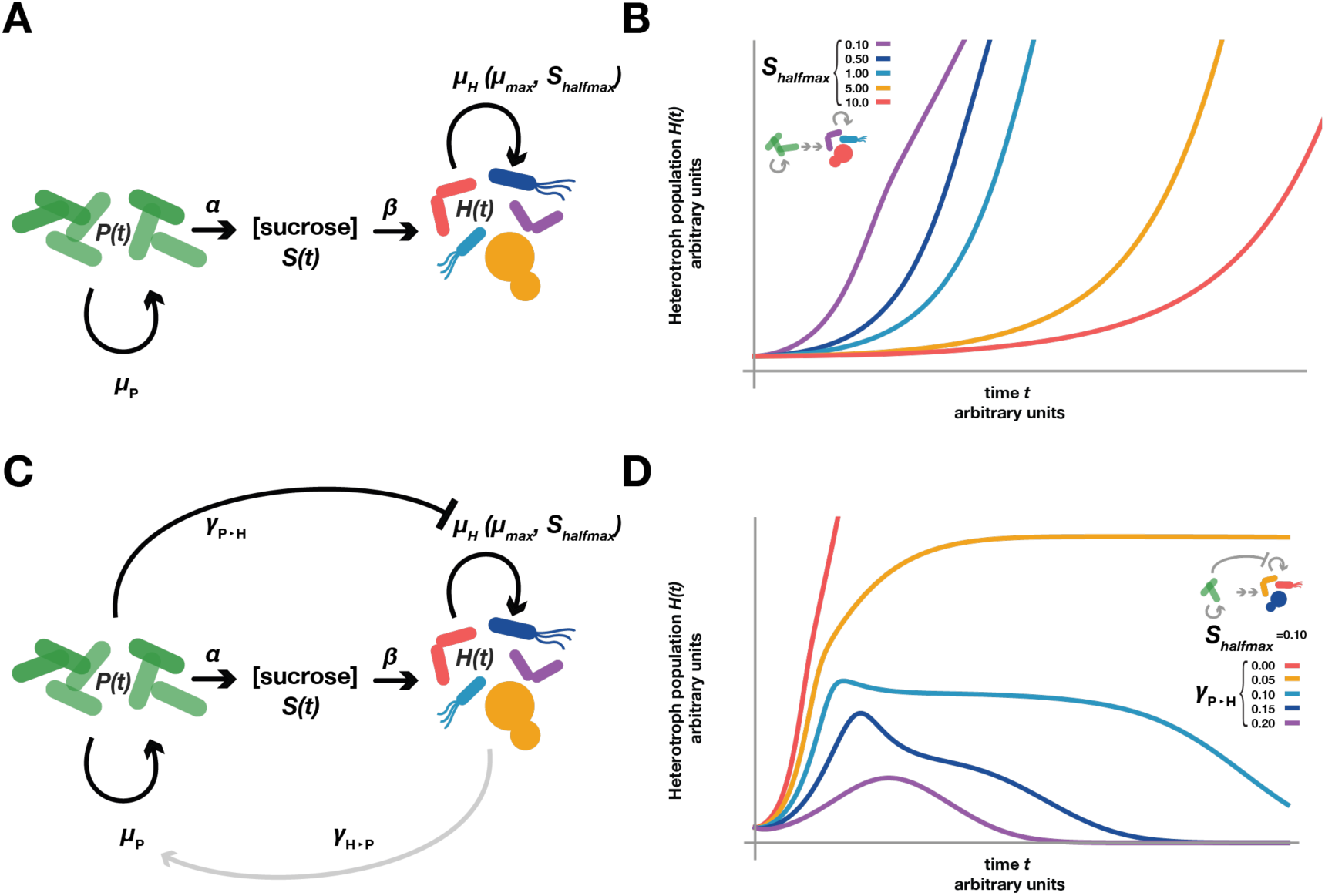
Mathematical Framework Describing Co-culture Interactions. (A, C) A mathematical framework summarizing consortia interactions we observe in this work is made up of: phototroph population (*P(t)*), phototroph growth rate (*μ*_*p*_), sucrose production rate by the phototroph (α), sucrose necessary for a heterotroph cell to double (*β*), the heterotroph population (*H(t)*), the heterotroph growth rate (*μ*_*H*_) which depends on the maximal growth rate when sucrose is not limiting (*μ*_*max*_) and the sucrose concentration at which the growth rate is half maximal (*S*_*halfmax*_), and the interaction terms describing the effects phototrophs haves on heterotrophs (*γ*_*P*⊣*H*_), and heterotrophs have phototrophs (γ_*H* →*P*_). (A, B) When interaction terms (*γ*_*P*⊣*H*_& γ_*H* →*P*_) are set to zero, and all other variables are held constant, decreases in *S*_*halfmax*_ increase growth and decrease time before the onset of heterotrophic growth. (C, D) Adding and tuning the *γ*_*P*⊣*H*_ interaction term stops the monotonically increasing heterotroph growth when other variables are held constant; at high enough values, the heterotrophic population can decrease and/or growth is entirely eliminated. Changes in γ_*H* →*P*_ (grey arrows in C, set to a value of ‘0’ in panel D) can also lead to feedback that increases the complexity of heterotrophic growth when other variables are held constant. As γ_*H* →*P*_ increases there is a more pronounced drop in heterotroph concentration after the initial rise.

We added variables to this framework to represent the additional observed interactions between phototrophs and heterotrophs (Fig. 6C): cyanobacterial inhibition of heterotrophic viability (Fig. 3A) and heterotrophic stimulation of cyanobacterial growth (Fig. 3E). The inhibitory effect of cyanobacteria on heterotroph, *γ*_*P*⊣*H*_, scales as a function of the phototroph density *P(t)*, and negatively influences the heterotroph population *H(t)*. The growth-promoting effect of heterotrophs on cyanobacterial growth, γ_*H* →*P*_, is represented as a constant that influences the cyanobacterial growth rate, μ*p*, as we have not shown this effect to be density-dependent.

Although limited to being a conceptual tool, this simplistic mathematical framework can recapitulate aspects of the complex interactions we observe in our experiments (Fig. 6). For example, when the interaction terms, *γ*_*P⊣H*_ and *γ*_*H→P*_, are set to 0 and all other parameters are held constant (details in Supplemental Material), increasing sucrose utilization (i.e. decreasing *S*_*halfmax*_) results in an earlier rise in heterotrophic population (Fig. 6A&B). This is consistent with data from *S. cerevisiae* co-cultures, as the W303 strain exhibits a high S_halfmax_ (Fig. 2C) and does not exhibit growth within co-culture (Fig. 2B) while the W303^Clump^ strain has a lower S_halfmax_ (Fig. 2C), and demonstrates growth within co-cultures (Fig. 2D).

More interestingly, the addition of the interaction term *γ*_*P⊣H*_ changes the shape of the heterotroph population function when all other parameters are constant; at higher values the model predicts an initial rise in heterotrophic density followed by a subsequent decline (Fig. 6D). This is consistent with instances in which the heterotroph species are shown to be fairly sensitive to the inhibitory effects of cyanobacterial metabolism in the light (demonstrated in Fig. 3A-D), and recapitulates experimental co-culture dynamics we observe with *S. cerevisiae* and *B. subtilis* (Figure 2A&D).

## Discussion

We show that *cscB*^+^ *S. elongatus* support diverse heterotrophic microbes in co-culture, demonstrating a flexible autotroph/heterotroph consortia platform. In comparison to other synthetic communities that have been constructed by cross-feeding [19–22, 36], sucrose is a metabolite that is naturally bioavailable to many microbes, and therefore the diversity of species with potential to be supported by *cscB*^+^ *S. elongatus* is broad. In constructing these consortia, we observed unforeseen interactions with common features shared across different heterotrophic species. Namely, we observe that light-driven processes of cyanobacteria have negative impacts on all tested heterotrophic species while, conversely, growth of all heterotrophic species simulates cyanobacterial growth. By taking measures to mitigate deleterious interactions, we are able to stabilize consortia over time and demonstrate consortia persistence in the face of fluctuations in light availability, population density, and composition. Finally, we show that these consortia can be functionalized to produce target compounds, where the end product is dictated by the heterotrophic partner.

Although the conceptual design of this platform is modular, we identify desirable features in a heterotrophic partner that could be important to maximize the stability and productivity of the coculture. A straightforward engineering target to improve stability and product output from cyanobacterial/heterotroph consortia is to enhance the efficiency of sucrose utilization in the heterotrophic partner. While the sucrose production of *cscB*^+^ *S. elongatus* is significant in comparison to traditional plant based feedstocks [10], it is produced continuously and the concentration remains low relative to standard laboratory media (frequently ~2% carbohydrate by volume). We see that use of a derivative yeast strain with mutations conferring enhanced utilization of sucrose (W303^Clump^ *S. cerevisiae*) greatly improves growth relative to the WT in co-culture with *cscB*^+^ *S. elongatus* (Fig. 2B&C), ultimately permitting stable co-culture over months in ePBRs (Fig. 4B, Additional File 1: Fig. S7&S8). Similarly, the deletion of the sucrose catabolism operon repressor cscR from the W strain of *E. coli* improved growth on low sucrose as expected (Fig. 2C) [27], and also greatly increased PHB production during week-long co-cultures experiments (Fig. 5D). However, there is still room for further improvement of sucrose uptake in Δ*cscR E. coli*, as evidenced by its insensitivity to *cscB*^+^ *S. elongatus* sucrose export during 48 hour batch cultures where sucrose concentrations remain < 1g/L (Fig. 2A, Additional File 1: Fig. S2). It is likely that the relative success of production and long-term stability of *ΔcscR E. coli* in co-culture with *cscB*^+^ *S. elongatus* (Fig. 4A, Additional File 1: Fig. S6) is due to an intrinsic capacity of this strain of *E. coli* to utilize other products cyanobacteria secrete that were not engineered as part of the consortia design.

Similarly, other emergent interactions that we consistently observe between *cscB*^+^ *S. elongatus* and all three heterotrophic species likely influence stability and productivity of a given co-culture pair. All three heterotrophs demonstrated decreased growth and viability when exposed to high densities of cyanobacteria in the light (Fig. 3A-D), suggesting that a cyanobacterially-derived product derived from active photosynthetic metabolism is the underlying cause. One possibility is that unavoidable byproducts of photosynthesis itself (e.g. oxygenation and/or reactive oxygen species) may be key [28]. We hypothesize that O_2_ production may be a large component of this toxicity effect due to the light dependence of oxygen evolution and the detrimental effect too much oxygen can have on cells [37–40]. Indeed, we observed that the addition of the herbicide DCMU (which stops oxygen evolution through inhibition of photosystem II – but does not prevent ROS formation or cyclic electron flow[41]) can rescue growth of *B. subtilis*, when co-cultured with dense cyanobacteria in the light (Additional File 1: Fig. S10). Consistent with this, many evolved consortia containing oxygenic phototrophs contain partners with strategies to mitigate reactive oxygen species, a normal byproduct of oxygenic photosynthesis [28, 42, 43]; similar strategies may be employed to further improve the stability of synthetic consortia. While cyanobacteria/heterotroph competition for specific media components could also contribute to the growth suppression we observe, we consider this to be a relatively minor factor: for example, heterotrophs readily grow in media conditioned by *S. elongatus* [9, 10].

The second emergent interaction that we observe in a species-independent manner is the stimulation of cyanobacterial growth in the presence of each heterotrophic species. These observations are analogous to stimulatory effects of heterotrophs on cyanobacteria and algae in numerous natural examples, such as interactions between *Prochlorococcus* and “helper heterotrophs” [42], or experiments where microalgae accumulate more biomass in the presence of other microbes than in isolation [44]. However, the generality of the positive effect we observe is somewhat surprising, as the complement of secreted bioproducts from *E. coli, B. subtilis*, and *S. cerevisiae* are likely different (and these species are not naturally prevalent in environments dominated by cyanobacteria; therefore there is no expectation of evolutionarily-preexisting pathways.). These results suggest that many different heterotrophs could boost yields of *S. elongatus*, a prospect with significant implications for scaled cyanobacterial cultivation. All aerobic heterotrophs can be expected to respire CO_2_, although the benefit of this additional inorganic carbon to *S. elongatus* is questionable as these experiments were conducted under CO_2_-enriched environments (2%). Instead, we hypothesize that these benefits arise from emergent cross-feeding of metabolite(s) and/or absorption of byproducts, a division of labor commonly observed in natural consortia; though this speculative interpretation requires additional investigation.

Our work represents an extension of multiple recent efforts in the design of synthetic microbial consortia: defined here as communities of two or more unrelated microbes that have been engineered to interact with one another through metabolic intermediates or molecular signals [45, 46]. For example, synthetic consortium engineering has been used as a bottom-up approach to gain insight into complex dynamics including population behavior, game theory, pattern formation, and cross-feeding [47–52]. The majority of described synthetic consortia involve the exchange of engineered signals (i.e. quorum sensing) or co-cultivation of complementary auxotrophs, frequently of the same microbial species [52–56]. These types of engineered interactions are relatively inflexible; because of the specialized metabolic signals such consortia designs are constrained to a limited number of species, confounding the identification of species-independent phenomena. Additionally, synthetic consortia are frequently fragile, functioning only for short time frames or requiring artificially structured environments [57, 58]. Continued advancement of synthetic consortia towards academic and industrial application will likely require platforms that address some of these concerns.

The *cscB*^+^ *S. elongatus* consortia system exhibits flexibility in that it supports the growth of three distinct “workhorse” model microbial organisms and these co-cultures can be stabilized over time and through perturbation. This flexibility of design allows for modular reconstruction of the consortia platform for a variety of light-driven applications: different heterotrophic organisms can be inoculated with *cscB^+^ S. elongatus* to reprogram the population for distinct functions. In this work, we show proof-of-principle that co-culture with cyanobacteria can drive the production of alpha-amylase from *B. subtilis* and PHB from *E. coli*, both commercially relevant products. Furthermore, while this manuscript was in preparation, it was reported that *cscB^+^ S. elongatus* can also support the growth and nitrogen fixation capacities of *Azotobacter vinelandii* [59]. To improve upon the productivity of these early designs and more completely capitalize upon the high sucrose productivities of *cscB^+^ S. elongatus*, it may be necessary to select or engineer heterotroph species with more favorable co-culture properties (e.g. superior sucrose uptake, resistance to hyperoxyia, or ability to use other cyanobacterial byproducts). Indeed, in related unpublished work, we have demonstrated that cosortia PHB specific productivities can be boosted by nearly three orders of magnitude by selection of a species with naturally favorable characteristics, *Halimonas bolievensis* (Weiss *et al.*, in preparation). Improved characteristics can also be engineered into a target heterotrophic strain, as we demonstrate here, and the platform provides a methodology to identify genetic determinants that would further improve consortia performance (e.g. laboratory evolution in long-term co-culture populations). Finally, the flexibility of this system allows for the determination of species-independent factors that promote cyanobacterial coexistence with other microbes, which may be useful for determining generalized interactions that underlie the formation and stabilization of natural cyanobacterial symbioses or more general phototroph/heterotroph symbioses [60–64].

## MATERIALS AND METHODS

### Strains, media, and axenic characterization

*S. elongatus* PCC 7942 (obtained from ATCC #33912) was engineered to secrete sucrose through the expression of the sucrose/proton symporter *cscB* [10]. *E. coli* W was obtained from ATCC (#9637) and the corresponding W Δ*cscR* strain was generously provided by Dr. Claudia Vicker’s laboratory [27]. *B. subtilis* 168 was obtained from ATCC (#23857) and *B. subtilis* 3610 *ΔsinI* was generously provided by the lab of Dr. Richard Losick [65]. The *ΔsinI* mutant strain of 3610 was used to minimize chained growth making CFU counts of the strain reproducible [65]. *S. cerevisiae* strains, WT W303 and W303^Clump^ (previously referred to as Ancestor and Recreated02 strains, respectively) were generously provided by the lab of Dr. Andrew Murray [23]. All strains are listed in Table 1.

**Table 1.** Strain and plasmid collection.

| **Strain** | **Origin** |
| --- | --- |
| *Synechococcus elongatus* PCC7942 | ATCC 33912 |
| *Synechococcus elongatus* trc-lac/cscB | [10] |
| *Bacillus subtilis* 168 | ATCC 23857 |
| *Bacillus subtilis* 3610 ΔsinI | [65] |
| *Escherichia coli* K-12 BW25113 | [68] |
| *Escherichia coli* W | ATCC 9637 |
| *Escherichia coli* W ΔcscR | [27] |
| *Saccharomyces cerevisiae* W303 | Ancestor strain; [23] |
| *Saccharomyces cerevisiae* W303<sup>Clump</sup> | Recreated02 strain; [23] |
| **Plasmid** |  |
| pAET41 | [33] |

*S. elongatus* was propagated in BG-11 (Sigma-Aldrich) plus 1g/L HEPES, pH 8 in constant light at 35 °C. Axenic cyanobacteria were checked for contamination via plating on rich media. *B. subtilis* and *E. coli* were propagated in LB-Miller (EMD Millipore) while *S. cerevisiae* was maintained in YEPD media (MP Biomedicals). *E. coli*, *B. subtilis*, and *S. cerevisiae* were struck from frozen stocks on rich media plates (LB for bacteria and YEPD for yeast). Co-culture media were optimized for either prokaryotes (^CoB^BG-11) or *S. cerevisiae* (^CoY^BG-11). ^CoB^BG-11 consists of BG-11 supplemented with 106mM NaCl, 4mM NH_4_Cl and 25mM HEPPSO, pH 8.3-KOH. Indole (100µM) was added to *B. subtilis* 168 co-cultures as indicated and in alpha-amylase experiments. ^CoY^BG-11 consists of BG-11 supplemented with 0.36g/L Yeast Nitrogen Base without amino acids (Sigma Aldrich), 106mM NaCl, 25mM HEPPSO, pH 8.3-KOH and 1mM KPO_3_. Solid co-culture plates were composed of ^CoB^BG-11 media with 1% autoclaved noble agar (BD Biosciences).

For characterization of *S. elongatus* growth and sucrose production, *S. elongatus* was cultured axenically in baffled flasks of ^CoB^BG-11 or ^CoY^BG-11 and allowed to acclimate for ≥ 12 hours. Then cultures were adjusted to 25mL with a final density of 0.5 OD_750_. IPTG (1mM) was added, as appropriate. This was the start of the experiment and is referred to as time 0. Cultures were monitored at 24 hour intervals by withdrawal of 1mL culture. OD_750_was measured via photospectrometer (ThermoScientific NonoDrop 2000c) and culture supernatant was analyzed for sucrose content via a colorimetric Glucose-Sucrose Assay (Megazyme).

To prepare heterotrophic strains, single colonies were picked into their respective rich media and grown until turbid at varying temperatures before co-culture (37 °C for *E. coli* and *B. subtilis*; 30 °C for *S. cerevisiae*). Cells were diluted into the appropriate co-culture media +2% sucrose to acclimate to coculture media, and maintained within log phase growth (OD_600_ < 0.70) before use in co-cultures. All acclimating cultures and co-cultures were grown at 35°C, 150rpm, 2% CO_2_, in light (PAR = ~80µmol with 15W Gro-Lux Sylvania fluorescent bulbs) within a Multitron Infors HT incubator. Heterotrophic growth was measured by inoculating rinsed cells at 0.01 OD_600_ (bacteria) or 0.05 OD_600_ (yeast) into fresh co-culture media at the indicated sucrose concentration. Data for growth rate was collected from 25mL flask cultures while 96-well plates with 1mL culture volumes were used to assay growth in a gradient of sucrose concentrations (.156mg/mL to 10mg/mL, Fig. 2C); OD_600_ of plates were read on a BioTek Synergy Neo plate reader.

### Batch co-cultivation & quantification

Flask co-cultures were completed in 25mL volumes in baffled flasks. Cyanobacteria and heterotrophs were acclimated to ^CoB^BG-11 or ^CoY^BG-11 media prior to inoculation into co-cultures. All co-cultures were grown at 35°C, 150rpm, 2% CO_2_, in light (15W; Gro-Lux; Sylvania) within a Multitron Infors HT incubator. 1mM IPTG was added when indicated. Growth in co-cultures was monitored every 12 hours: *S. elongatus* was measured by the count of gated red-fluorescent events on a quantitative flow cytometer (BD Accuri); heterotrophs were assayed by plating dilution series on rich media to count colony forming units (CFU). Estimates of W303^Clump^ cell number were derived by counting CFUs, but numbers were adjusted for the ~6.6cells/clump as previously reported [23], and as confirmed under our culture conditions. For dilution experiments, co-cultures containing *E. coli* or *B. subtilis* were grown for 24 hours before 10 or 100 fold dilutions.

### Heterotroph exposure to variable cyanobacteria densities

*B. subtilis* and *E. coli* were recovered from rich media as above, washed in ^CoB^BG-11 and inoculated at an OD_600_ of .01 in ^CoB^BG-11 media + 2% sucrose with cyanobacteria at different densities (OD_750_ 0, 0.5, 1, and 2). *S. cerevisiae* was treated identically except they were inoculated at ~3x10^5^cells/mL (OD_750_ = 0.03) and ^CoY^BG-11 was used. These samples were split into two 36-well plates and incubated and exposed to either constant light or dark conditions while maintaining the other growth parameters. Heterotroph counts were determined by plating on rich media for colony counts as above after initial setup (time 0) and after 12 hours of culture. Ratios of the viable cell counts from the light vs. dark cultures or log_10_ of these ratios after 12 hours are reported.

### Structured growth perturbation

To test the ability of co-cultures to withstand environmental perturbation, flask co-cultures were inoculated and grown as previously described for 24 hours before plating of 100µL on solid co-culture Petri dishes. After five days, uneven lawns of heterotrophs and cyanobacteria arose. Cells were picked from these plates into 96-well plates and allowed to grow for 2-5 additional days. Any well that demonstrated cyanobacterial growth (as judged visually by green appearance) at the end of 48 hours was spotted on rich media to determine the presence or absence of heterotrophic symbionts. Solid culture and 96-well plate growth was completed at 35°C, 0rpm, 2% CO_2_, in constant light (15W; Gro-Lux; Sylvania) within a Multitron Infors HT incubator.

### Heterotroph spotting on cyanobacterial lawns

Lawns of *cscB*^+^ cyanobacteria were achieved via spreading of 250µL of *cscB*^*+*^ cyanobacteria (OD_750_ 0.5) on solid co-culture plates with or without 1mM IPTG. After the cyanobacteria had absorbed on to the plate (>3 hours in the dark), 3µL drops of heterotrophs were spotted on to the lawns. Heterotrophs had been previously grown up in rich media and washed three times to remove any media components before spotting. Media blanks and boiled cells were spotted as negative controls. Plates were then grown at 35°C, 2% CO_2_, in constant light (15W; Gro-Lux; Sylvania) within a Multitron Infors HT incubator.

### Long-term continuous co-cultivation

Long-term co-cultures were incubated in Phenometrics Environmental Photo-Bioreactors [30] with 150mL liquid volumes of a mix of *cscB^+^ S. elongatus* with either *S. cerevisiae* W303^Clump^ or *E. coli* W Δ*cscR* in the appropriate co-culture BG-11 media + 1mM IPTG. Reactors were seeded with ~1x10^8^cells/mL of *S. elongatus* (OD_750_ = 0.5) and a final concentration of heterotroph equivalent to ~1x10^6^cells/mL *S. cerevisiae* W303^Clump^ (final OD_600_ ~0.1) or ~5x10^7^cells/mL *E. coli* W Δ*cscR* (OD_600_ ~0.05). Light was provided by onboard white, high-power LEDs (400µmol m^2^ s^2^) continuously for *E. coli* W Δ*cscR* cultures, and with a 16:8 light:dark photoperiod for *S. cerevisiae* W303^Clump^ co-cultures. The total density of co-cultures was monitored by onboard infrared diodes, following a brief (3-12 hour) acclimation period where the time-averaged optical density was allowed to settle to a fixed point following culture initiation. This measurement was used to control attached peristaltic pumps that eject fresh media to maintain the set target OD as previously described [30]. Co-culture temperature was maintained at 30°C by a heated jacket; cells were agitated continuously by a stirbar. Daily, ~2mL of co-culture volume was withdrawn and cyanobacterial and heterotrophic cell counts determined by flow cytometry and plating, respectively (as described above).

### Alpha-amylase production and quantification

For the production of alpha-amylase, co-cultures of *cscB*^*+*^ *S. elongatus* and *B. subtilis* strain 168 were completed in 8mL volumes of ^CoB^BG-11 supplemented with 100µM indole in 6 well dishes. When specified, cyanobacteria were present (OD_750_ = 0.5) with or without 1mM IPTG. Control cultures did not contain cyanobacteria. Alpha-amylase production was measured after 24 hours of culture at 35°C, 0rpm, 2% CO_2_, in constant light (15W; Gro-Lux; Sylvania) within a Multitron Infors HT incubator. Alpha-amylase activity in supernatants was measured immediately after pelleting of cultures with the EnzChek Ultra Amylase Assay Kit, Molecular Probes Life Technologies using the manufacturer’s protocol. Western blots confirmed presence of alpha-amylase in supernatants after addition of NuPAGE LDS sample buffer (Invitrogen) followed by 10 minutes at 100°C. Protein (10µL) was run on NuPage 4-12% Bis-Tris gels (Life Technologies) for in MES SDS running buffer for 50 minutes at 185V. The iBlot 2 Dry Blot System (ThermoScientific) was used to transfer protein to nitrocellulose membranes (iBlot 2 NC Regular Transfer Stacks). Anti-alpha amylase antibodies (polyclonal rabbit; LS-C147316; LifeSpan BioSciences; 1:3,000 dilution) were used as the primary antibody followed by peroxidase-conjugated donkey anti-rabbit antibodies (AffiniPure 711-035-152 lot 92319; Jackson ImmunoResearch; 1:5,000 dilution) as the secondary antibody. The western blot was visualized via Western Lightning^®^ Plus-ECL, Enhanced Chemiluminescence Substrate (PerkinElmer, ProteinSimple FluorChem M). Purified alpha-amylase (Sigma Aldrich) was used as a control in all assays.

### PHB production & quantification

*E. coli* strains were transformed with pAET41 (Table 1) before use in co-cultures for production [33]. Co-cultures were set up as previously described in 25mL flasks. After one week of growth, the entire culture was spun down, frozen, and stored at −80°C until PHB content was quantified. PHB content was quantified by standard methods [66, 67]. Briefly: cell pellets were digested with concentrated H_2_SO_4_ at 90°C for 60 min. The digestion solution was diluted with H_2_O by 500 times and passed through 0.2µm filter. The solutions were subsequently analyzed by a high performance liquid chromatography (HPLC, Agilent HPLC 1200) equipped with Aminex HPX-87H column and UV absorption detector [67]. The volume of each sample injection was 100μL. The mobile phase was 2.5mM H_2_SO_4_ aqueous solution, with a flow rate of 0.5mL/min for 60min. 5mM sodium acetate (Sigma Aldrich) was added as an internal standard. The concentrations of PHB were determined by comparing the peak area with that in standard curves from 0.1 to 30mM.

### Mathematical framework, statistics, and figures

All equations were modeled in Mathematica (Wolfram Research, Inc., Mathematica, Version 11.0). Statistics were completed in GraphPad Prism version 7, GraphPad Software, La Jolla California USA, www.graphpad.com

## LIST OF ABBREVIATIONS

**PHB -** polyhydroxybutyrate

**IPTG -** isopropyl β-D-1-thiogalactopyranoside

**CFU -** colony forming unit

**WT -** wild type

**ePBR -** environmental Photobioreactors

## Footnotes

**AUTHORS’ CONTRIBUTIONS:** DD and SH conceived and designed experiments. SH, DD, and LY performed experiments. DD, SH, and PS wrote the manuscript. All authors read and edited the manuscript.

## ACKNOWLEDGEMENTS

Special thanks to Chong Liu and the Nocera laboratory for quantifying PHB accumulation and Charleston Noble from the Nowak laboratory for helping us explore mathematical frameworks of our communities. Additionally, we are grateful to the Betenbaugh, Guarnieri, and Zengler groups for their conversations and collaboration. We must also thank the members of the Ducat (especially Jingcheng Huang, Brad Abramson, Eric Young, Josh MacCready, Derek Fedeson, Taylor Weiss), and Silver (especially John Oliver, Brendan Colon, Marika Ziesack) labs as well as Jenna Chandler, Igor Vieira, Rebecca Ward, and Henry Wettersten for their critical reviews of the manuscript.

## REFERENCES

1. Hays SG, Ducat DC: Engineering cyanobacteria as photosynthetic feedstock factories. Photosynth Res 2015, 123:285–95.

2. Du W, Liang F, Duan Y, Tan X, Lu X: Exploring the photosynthetic production capacity of sucrose by cyanobacteria. Metab Eng 2013, 19:17–25.

3. Song K, Tan X, Liang Y, Lu X: The potential of Synechococcus elongatus UTEX 2973 for sugar feedstock production. Appl Microbiol Biotechnol 2016, 100:7865–7875.

4. Möllers KB, Cannella D, Jørgensen H, Frigaard N-U: Cyanobacterial biomass as carbohydrate and nutrient feedstock for bioethanol production by yeast fermentation. Biotechnol Biofuels 2014, 7:64.

5. Aikawa S, Joseph A, Yamada R, Izumi Y, Yamagishi T, Matsuda F, Kawai H, Chang J-S, Hasunuma T, Kondo A, Ducat DC, Way JC, Silver PA, Lopez PJ, Desclés J, Allen AE, Bowler C, Rosenberg JN, Oyler GA, Wilkinson L, Betenbaugh MJ, Melis A, Dismukes GC, Carrieri D, Bennette N, Ananyev GM, Posewitz MC, Harun R, Jason WSY, Cherrington T, et al.: Direct conversion of Spirulina to ethanol without pretreatment or enzymatic hydrolysis processes. Energy Environ Sci 2013, 6:1844.

6. Markou G, Angelidaki I, Georgakakis D: Carbohydrate-enriched cyanobacterial biomass as feedstock for bio-methane production through anaerobic digestion. Fuel 2013, 111:872–879.

7. Duan Y, Luo Q, Liang F, Lu X: Sucrose secreted by the engineered cyanobacterium and its fermentability. J Ocean Univ China 2016, 15:890–896.

8. Xu Y, Guerra LT, Li Z, Ludwig M, Dismukes GC, Bryant DA: Altered carbohydrate metabolism in glycogen synthase mutants of Synechococcus sp. strain PCC 7002: Cell factories for soluble sugars. Metab Eng 2013, 16:56–67.

9. Niederholtmeyer H, Wolfstädter BT, Savage DF, Silver PA, Way JC: Engineering cyanobacteria to synthesize and export hydrophilic products. Appl Environ Microbiol 2010, 76:3462–6.

10. Ducat DC, Avelar-Rivas JA, Way JC, Silver PA: Rerouting carbon flux to enhance photosynthetic productivity. Appl Environ Microbiol 2012, 78:2660–8.

11. Xu Y, Guerra LT, Li Z, Ludwig M, Dismukes GC, Bryant DA: Altered carbohydrate metabolism in glycogen synthase mutants of Synechococcus sp. strain PCC 7002: Cell factories for soluble sugars. Metab Eng 2013, 16:56–67.

12. Aikens J, Turner RJ: Method of producing a fermentable sugar. 2013.

13. Chisti Y: Constraints to commercialization of algal fuels. J Biotechnol 2013, 167:201–14.

14. Bux F, Chisti Y (Eds): Algae Biotechnology. Cham: Springer International Publishing; 2016. [Green Energy and Technology]

15. Singh J, Gu S: Commercialization potential of microalgae for biofuels production. Renew Sustain Energy Rev 2010, 14:2596–2610.

16. Ortiz-Marquez JCF, Do Nascimento M, Zehr JP, Curatti L: Genetic engineering of multispecies microbial cell factories as an alternative for bioenergy production. Trends Biotechnol 2013, 31:521–529.

17. Klähn S, Hagemann M: Compatible solute biosynthesis in cyanobacteria. Environ Microbiol 2011, 13:551–562.

18. Bockmann J, Heuel H, Lengeler JW: Characterization of a chromosomally encoded, non-PTS metabolic pathway for sucrose utilization in Escherichia coli EC3132. MGG Mol Gen Genet 1992, 235:22–32.

19. Song H-S, Renslow RS, Fredrickson JK, Lindemann SR: Integrating Ecological and Engineering Concepts of Resilience in Microbial Communities. Front Microbiol 2015, 6:1298.

20. Kim JH, Boedicker JQ, Choi JW, Ismagilov RF: Defined spatial structure stabilizes a synthetic multispecies bacterial community. Proc Natl Acad Sci 2008, 105:18188–18193.

21. Wintermute EH, Silver PA: Dynamics in the mixed microbial concourse. Genes Dev 2010, 24:2603–14.

22. Mee MT, Collins JJ, Church GM, Wang HH: Syntrophic exchange in synthetic microbial communities. Proc Natl Acad Sci U S A 2014, 111:E2149x–56.

23. Koschwanez JH, Foster KR, Murray AW: Improved use of a public good selects for the evolution of undifferentiated multicellularity. Elife 2013, 2013:1–27.

24. Koschwanez JH, Foster KR, Murray AW, Keller L: Sucrose Utilization in Budding Yeast as a Model for the Origin of Undifferentiated Multicellularity. 2011.

25. Archer CT, Kim JF, Jeong H, Park J, Vickers CE, Lee S, Nielsen LK: The genome sequence of E. coli W (ATCC 9637): comparative genome analysis and an improved genome-scale reconstruction of E. coli. BMC Genomics 2011, 12:9.

26. Sabri S, Nielsen LK, Vickers CE: Molecular control of sucrose utilization in Escherichia coli W, an efficient sucrose-utilizing strain. Appl Environ Microbiol 2013, 79:478–487.

27. Arifin Y, Sabri S, Sugiarto H, Krömer JO, Vickers CE, Nielsen LK: Deletion of cscR in Escherichia coli W improves growth and poly-3-hydroxybutyrate (PHB) production from sucrose in fed batch culture. J Biotechnol 2011, 156:275–8.

28. Villa F, Pitts B, Lauchnor E, Cappitelli F, Stewart PS: Development of a laboratory model of a phototroph-heterotroph mixed-species biofilm at the stone/air interface. Front Microbiol 2015, 6(NOV):1–14.

29. Rossi F, De Philippis R: Role of Cyanobacterial Exopolysaccharides in Phototrophic Biofilms and in Complex Microbial Mats. Life 2015, 5:1218–1238.

30. Lucker BF, Hall CC, Zegarac R, Kramer DM: The environmental photobioreactor (ePBR): An algal culturing platform for simulating dynamic natural environments. Algal Res 2014, 6(PB):242–249.

31. van Dijl JM, Hecker M: Bacillus subtilis: from soil bacterium to super-secreting cell factory. Microb Cell Fact 2013, 12:3.

32. Green DM, Colarusso LJ: The physical and genetic characterization of a transformable enzyme: Bacillus subtilis αamylase. Biochim Biophys Acta - Spec Sect Enzymol Subj 1964, 89:277–290.

33. Peoples OP, Sinskey AJ: Poly-??-hydroxybutyrate (PHB) biosynthesis in Alcaligenes eutrophus H16. Identification and characterization of the PHB polymerase gene (phbC). J Biol Chem 1989, 264:15298–15303.

34. Monod J: Recherches Sur La Croissance Des Cultures Bactériennes,. Paris: Hermann & cie; 1942.

35. Smith HL: Bacterial Growth.

36. Song H, Ding M-Z, Jia X-Q, Ma Q, Yuan Y-J: Synthetic microbial consortia: from systematic analysis to construction and applications. Chem Soc Rev Chem Soc Rev 2014, 6954:6954–6981.

37. Cross JB, Currier RP, Torraco DJ, Vanderberg LA, Wagner GL, Gladen PD: Killing of bacillus spores by aqueous dissolved oxygen, ascorbic acid, and copper ions. Appl Environ Microbiol 2003, 69:2245–52.

38. Outten EC, Falk RL, Culotta VC: Cellular factors required for protection from hyperoxia toxicity in Saccharomyces cerevisiae. Biochem J 2005, 388:93–101.

39. Kolesky DB, Truby RL, Gladman a. S, Busbee T a., Homan K a., Lewis J a.: 3D bioprinting of vascularized, heterogeneous cell-laden tissue constructs. Adv Mater 2014, 26:3124–3130.

40. Baez A, Shiloach J: Effect of elevated oxygen concentration on bacteria, yeasts, and cells propagated for production of biological compounds. Microb Cell Fact 2014, 13:181.

41. Das PK, Bagchi SN: Bentazone and bromoxynil induce H+ and H2O2 accumulation, and inhibit photosynthetic O2 evolution in Synechococcous elongatus PCC7942. Pestic Biochem Physiol 2010, 97:256–261.

42. Morris JJ, Kirkegaard R, Szul MJ, Johnson ZI, Zinser ER: Facilitation of robust growth of Prochlorococcus colonies and dilute liquid cultures by “helper” heterotrophic bacteria. Appl Environ Microbiol 2008, 74:4530–4534.

43. Beliaev AS, Romine MF, Serres M, Bernstein HC, Linggi BE, Markillie LM, Isern NG, Chrisler WB, Kucek LA, Hill EA, Pinchuk GE, Bryant DA, Wiley S, Fredrickson JK, Konopka A: Inference of interactions in cyanobacterial–heterotrophic co-cultures via transcriptome sequencing. ISME J 2014, 869:2243–2255.

44. Do Nascimento M, Dublan M de los A, Ortiz-Marquez JCF, Curatti L: High lipid productivity of an Ankistrodesmus-Rhizobium artificial consortium. Bioresour Technol 2013, 146:400–7.

45. Bernstein HC, Carlson RP: Microbial Consortia Engineering for Cellular Factories: in vitro to in silico systems. Comput Struct Biotechnol J 2012, 3:e201210017.

46. Hays SG, Patrick WG, Ziesack M, Oxman N, Silver PA: Better together: Engineering and application of microbial symbioses. Curr Opin Biotechnol 2015, 36:40–49.

47. Balagaddé FK, Song H, Ozaki J, Collins CH, Barnet M, Arnold FH, Quake SR, You L: A synthetic Escherichia coli predator-prey ecosystem. Mol Syst Biol 2008, 4:187.

48. Yurtsev EA, Conwill A, Gore J: Oscillatory dynamics in a bacterial cross-protection mutualism. Proc Natl Acad Sci U S A 2016, 113:6236–41.

49. Gore J, Youk H, van Oudenaarden A: Snowdrift game dynamics and facultative cheating in yeast. Nature 2009, 459:253–6.

50. Chen Y, Kim JK, Hirning AJ, Josić K, Bennett MR: Emergent genetic oscillations in a synthetic microbial consortium. Science (80-) 2015, 349.

51. Basu S, Gerchman Y, Collins CH, Arnold FH, Weiss R: A synthetic multicellular system for programmed pattern formation. Nature 2005.

52. Shou W, Ram S, Vilar JMG: Synthetic cooperation in engineered yeast populations. Proc Natl Acad Sci U S A 2007.

53. Tamsir A, Tabor JJ, Voigt CA: Robust multicellular computing using genetically encoded NOR gates and chemical “wires.” Nature 2011, 469:212–215.

54. Scott SR, Hasty J: Quorum Sensing Communication Modules for Microbial Consortia. 2016.

55. Wintermute EH, Silver PA: Emergent cooperation in microbial metabolism. Mol Syst Biol 2010, 6:407.

56. Mee MT, Collins JJ, Church GM, Wang HH: Syntrophic exchange in synthetic microbial communities. Proc Natl Acad Sci 2014, 20.

57. Song H-S, Renslow RS, Fredrickson JK, Lindemann SR: Integrating Ecological and Engineering Concepts of Resilience in Microbial Communities. Front Microbiol 2015, 6:1298.

58. Kim HJ, Boedicker JQ, Choi JW, Ismagilov RF: Defined spatial structure stabilizes a synthetic multispecies bacterial community. Proc Natl Acad Sci U S A 2008, 105:18188–18193.

59. Smith MJ, Francis MB: A Designed A. vinelandii-S. elongatus Coculture for Chemical Photoproduction from Air, Water, Phosphate, and Trace Metals. ACS Synth Biol 2016.

60. Rai AN, Bergman B, Rasmussen U: Cyanobacteria in Symbiosis. Dordrecht: Springer Netherlands; 2002.

61. Martínez-Pérez C, Mohr W, Löscher CR, Dekaezemacker J, Littmann S, Yilmaz P, Lehnen N, Fuchs BM, Lavik G, Schmitz RA, LaRoche J, Kuypers MMM: The small unicellular diazotrophic symbiont, UCYN-A, is a key player in the marine nitrogen cycle. Nat Microbiol 2016, 1:16163.

62. Aschenbrenner IA, Cernava T, Berg G, Grube M: Understanding Microbial Multi-Species Symbioses. Front Microbiol 2016, 7:180.

63. Hom EFY, Murray AW: Plant-fungal ecology. Niche engineering demonstrates a latent capacity for fungal-algal mutualism. Science 2014, 345:94–8.

64. Ramanan R, Kim B-H, Cho D-H, Oh H-M, Kim H-S: Algae-bacteria interactions: Evolution, ecology and emerging applications. Biotechnol Adv, 34:14–29.

65. Kearns DB, Chu F, Branda SS, Kolter R, Losick R: A master regulator for biofilm formation by Bacillus subtilis. Mol Microbiol 2005, 55:739–49.

66. Torella JP, Gagliardi CJ, Chen JS, Bediako DK, Colón B, Way JC, Silver PA, Nocera DG: Efficient solar-to-fuels production from a hybrid microbial–water-splitting catalyst system. Proc Natl Acad Sci 2015, 112:2337–2342.

67. Liu C, Gallagher JJ, Sakimoto KK, Nichols EM, Chang CJ, Chang MCY, Yang P: Nanowire–Bacteria Hybrids for Unassisted Solar Carbon Dioxide Fixation to Value-Added Chemicals. 2015.

68. Datsenko KA, Wanner BL: One-step inactivation of chromosomal genes in Escherichia coli K-12 using PCR products. Proc Natl Acad Sci U S A 2000, 97:6640–5.

